# Oxycodone Self-Administration in Female Rats is Enhanced by Δ^9^-tetrahydrocannabinol, but not by Cannabidiol, in a Progressive Ratio Procedure

**DOI:** 10.1101/2023.10.26.564282

**Authors:** Jacques D. Nguyen, Yanabel Grant, Celine Yang, Arnold Gutierrez, Michael A. Taffe

## Abstract

Epidemiological evidence suggests that the legalization of cannabis may reduce opioid-related harms. Preclinical evidence of neuropharmacological interactions of endogenous cannabinoid and opioid systems prompts further investigation of cannabinoids as potential therapeutics for the non-medical use of opioids. In these studies female rats, previously trained to self-administer oxycodone (0.15 mg/kg/infusion) intravenously in 6 h sessions, were allowed to self-administer oxycodone after exposure to cannabidiol (CBD) and Δ^9^-tetrahydrocannabinol (THC) by vapor inhalation and THC by injection (5.0-20 mg/kg, i.p.). Self-administration was characterized under Progressive Ratio (PR) and Fixed Ratio (FR) 1 schedules of reinforcement in 3 h sessions. THC decreased IVSA of oxycodone in a FR procedure but increased reward seeking in a PR procedure. CBD decreased the IVSA of oxycodone in the FR but not the PR procedure. The results are consistent with an anti-reward effect of CBD but suggest THC acts to increase the reinforcing efficacy of oxycodone in this procedure.

## 1 Introduction

Reduction of opioid use concomitant with cannabis use (Kim et al., 2016; Nguyen et al., 2023; Pardo, 2017; Piper et al., 2017; Shi, 2017) may arise because cannabis acts therapeutically to reduce the motivation to seek opioids, because it is substituting for the opioid or because it is enhancing the impact of a unit dose of the opioid. Cannabis smoking slightly increased the abuse-related subjective effects of oxycodone in a human laboratory study (Cooper et al., 2018), but *self-administration* was not directly assessed. It is difficult to experimentally determine if cannabis reduces opioid use in humans, but animal models can assist in distinguishing pro-reward from anti-abuse effects.

A recent investigation in a nonhuman primate model concluded there was no impact of cannabinoids on opioid rewarding efficacy (Maguire and France, 2020) however that study used a full agonist cannabinoid and the self-administration of a very short acting opioid, remifentanil, with the explicit intent that animals could not reach a satiety point for remifentanil under the experimental conditions. Use of a drug-food choice procedure (Carey et al., 2023) may suggest that the cannabinoid would have no effect on relative drug vs food motivation, but this potential interaction is not well established by the model. Δ^9^-tetrahydrocannabinol (THC) reinstates beer-and sucrose-seeking behavior in rats, for example (McGregor et al., 2005). THC has also been shown to *increase* the reinforcing efficacy of cocaine in a monkey model, but THC is not itself readily self-administered intravenously by rhesus monkeys (John et al., 2017; Mansbach et al., 1994) or by rats (Takahashi and Singer, 1979); for review see Justinova et alia (Justinova et al., 2005).

In our recent work we concluded that parallel shifts in the dose-response function for THC treatment across different unit doses of oxycodone, combined with an increase in PR breakpoint, is most compatible with the interpretation that THC enhances the rewarding value of a unit dose of self-administered oxycodone (Nguyen et al., 2019).

In animal models it is possible that a generally sedating effect of cannabinoids may disrupt behavioral responding non-specifically, effects that are misinterpreted as being specific to the motivation to seek drug effects in a self-administration procedure. Caution should be used when interpreting the impact of full-agonist compounds that reduce response rate where THC does not do so. This is also a reason why demonstration of increased behavioral responding in a PR procedure associated with THC is critical additional support for the interpretation of enhanced reinforcer efficacy.

Although Δ^9^-tetrahydrocannabinol is undoubtedly the constituent of cannabis that drives most use, for both recreational and therapeutic purposes, additional constituents may also produce significant effects. Cannabidiol (CBD) reduces alcohol or cocaine seeking behavior in rats when applied via a dermal gel (Gonzalez-Cuevas et al., 2018), as well as cocaine (Galaj et al., 2020) or methamphetamine (Hay et al., 2018) seeking in rats and alcohol seeking in rats (Tringali et al., 2023) and mice (Viudez-Martinez et al., 2017) when CBD is delivered by i.p. injection. CBD by intraventricular injection, speeds extinction, and attenuates methamphetamine cued re-instatement, of methamphetamine conditioned place preference (Mirmohammadi et al., 2022). There does not appear to be any evidence whether CBD can alter the rewarding value of opioids. A recent review of studies involving human substance users found most studies are of low quality and only hinted at a general beneficial effect of CBD on opioid craving and anxiety (Lo et al., 2023), e.g., (Suzuki et al., 2022); also see (Fernandes et al., 2023; Paulus et al., 2022). The effect may be route and/or species specific since effects of CBD in rats have only variably been demonstrated with parenteral injection (Boggs et al., 2018). There is no effect of oral CBD on sustained attention in rats (Moore et al., 2023). Furthermore, CBD vapor inhalation decreases rat body temperature in a partially serotonin 1A receptor-dependent manner (Javadi-Paydar et al., 2019), but no effects on body temperature are observed after CBD injection (Taffe et al., 2015). The impact of CBD may also be species dependent since oral cannabidiol did not affect the self-administration of alcohol in baboons (Moore et al., 2023).

This study was therefore conducted to determine if treatment with THC or CBD by vapor inhalation alters the intravenous self-administration of oxycodone in a rat model.

## 2 METHODS

### 2.1 Subjects

Subjects were female Wistar rats (Charles River), aged 10-11 weeks on arrival in the laboratory in a single cohort of 20. The vivarium was kept on a 12:12 hour reversed light-dark cycle, and behavior studies were conducted during the vivarium dark period. The animals were pair housed and food and water were provided ad libitum in the home cage and in the self-administration chambers. Experimental procedures were conducted in accordance with protocols approved by the Institutional Animal Care and Use Committee of the University of California, San Diego and consistent with recommendations in the NIH Guide (Garber et al., 2011).

#### Subject dropout

Two animals were lost due to mechanical failure of vivarium equipment prior to the start of intravenous self-administration (IVSA) training. Three other animals were lost to the study prior to the start due to catheter/implantation complications (cagemate chewed port off, failure to recover over three days, difficulty implanting catheter). One animal was discontinued for self-injurious behavior (paw chewing) after Session 27. Five animals were discontinued due to loss of catheter port integrity after Sessions 22, 31, 53, 54, and 64, respectively. The nine at the end of the study included 5 original THC group animals and 4 original Vehicle group animals.

### 2.2 Drugs

Vapor inhalation was conducted as previously described (Javadi-Paydar et al., 2019; Taffe et al., 2020) with the CBD or THC dissolved in propylene glycol (PG). In brief, commercially available e-cigarette cannisters (SMOK Baby Beast Brother TFV8 sub-ohm tanks equipped with V8 X-Baby M2 0.25 ohm coils) were triggered (La Jolla Alcohol Research, Inc Model SSV-3 or SVS-200 set to 58 watts) to fill an inhalation chamber (152 mm W X 178 mm H X 330 mm L) with vapor every five minutes over a 30 minute session. Animals were exposed in groups of 2-3 per vapor session.

### 2.3 Intravenous Self-Administration Acquisition

Female rats were implanted with intravenous catheters and trained to self-administer oxycodone (0.15 mg/kg/infusion) on a Fixed Ratio (FR) 1 schedule of reinforcement using methods previously described (Nguyen et al., 2019; Nguyen et al., 2021; Nguyen et al., 2018; Nguyen et al., 2017). In this study, separate groups were initially injected with the vehicle (N=7) or with THC (N=8) 30 minutes before each 6 h self-administration session; treatments were the same for housing pairs. The first 10 sessions were conducted on sequential days and the THC dose was 5 mg/kg, i.p. for the first 10 sessions, 10 mg/kg, i.p. for Sessions 11-14 and 20 mg/kg, i.p. for Sessions 15-19. IVSA was suspended for a week between Sessions 19 and 20 to permit assessment of the acute effects of THC (5, 10, 20 mg/kg, i.p.) on rectal temperature and nociception assays. For Sessions 22-28, the original Vehicle group was injected with 5 mg/kg, THC, i.p., prior to the IVSA sessions and the original THC group was injected with the vehicle. The response requirement was incremented to FR5 for Sessions 30-32.

### 2.4 Experiments

#### 2.4.1 Progressive Ratio

On sessions 33-37 animals were tested on a Progressive Ratio (PR) procedure with 0.15 mg/kg/infusion oxycodone available. In the PR paradigm, the required response ratio was increased after each reinforcer delivery, within a session (Hodos, 1961; Segal and Mandell, 1974) as determined by the following equation (rounded to the nearest integer): Response Ratio=5e^(injection number\**j*)–5 (Richardson and Roberts, 1996). In this study the *j* value was set to 0.2. Sessions were a maximum of 3 h in duration. Animals were injected with either Veh or THC (5 mg/kg, i.p.) before the 34th and 35th sessions in a counter-balanced order. The 36th and 37th sessions were conducted under PR, without any pre-session treatment. The per-infusion dose of oxycodone was changed to 0.06 mg/kg/inf for Sessions 38-41 and the rats were injected with either Veh or THC (5 mg/kg, i.p.) before the 39th and 40th sessions in a counter-balanced order. For analysis, the day following the injection days was used as the no-injection comparison.

#### 2.4.2 FR1

For Sessions 42-64, rats were permitted to self-administer oxycodone in 3 h sessions under a FR1 contingency. In Sessions 43-45 the oxycodone dose (0, 0.06, 0.15 mg/kg/infusion) was varied in a counter-balanced order.

The study next evaluated injection of 0, 5, 10 and 20 mg/kg of THC, i.p., prior to oxycodone (0.06 mg/kg/infusion) self-administration sessions, assessed in a counter-balanced order in Sessions 48-51.

IVSA was then suspended for a week between Sessions 53 and 54 to permit assessment of the acute effects of THC (5, 10, 20 mg/kg, i.p.) on rectal temperature and nociception assays.

Self-administration of oxycodone was returned to the original training dose (0.15 mg/kg/infusion) in 3 h sessions (FR1) for Sessions 56-64. Rats were exposed to cannabidiol (CBD) vapor (400 mg/mL in the PG) or vehicle (PG) vapor for 30 minutes immediately prior to IVSA sessions in a counter balanced order for Sessions 59-60. Two additional animals were discontinued due to catheter failure, thus N=10 (N=4 from original Veh group). Vapor inhalation was selected because we have previously reported effects of CBD on body temperature via this route (Javadi-Paydar et al., 2019; Javadi-Paydar et al., 2018) that were not observed following CBD injection of 30-60 mg/kg, i.p. (Taffe et al., 2015). Rats were next exposed to Air, vehicle (PG) vapor, or THC vapor (100 mg/mL in the PG) for 30 minutes immediately prior to oxycodone (0.15 mg/kg/infusion) IVSA sessions in a counter balanced order for Sessions 62-64.

**Table 1:** Summary of experiments by cumulative self-administration session.

| **IVSA Session** | **34-35,**<br>**39-40** | **43-45** | **48-51** | **59-60** | **62-64** | **66-68** | **71-73** | **75-77** | **86-88** |
| --- | --- | --- | --- | --- | --- | --- | --- | --- | --- |
| **Schedule of Reinforcement** | **PR** | **FR1** | **FR1** | **FR1** | **FR1** | **PR** | **PR** | **PR** | **FR1** |
| Oxycodone (mg/kg/infusion) | 0.06, 0.15 | 0.00, 0.06, 0.15 | 0.06 | 0.15 | 0.15 | 0.15 | 0.30 | 0.06, 0.15, 0.30 | 0.06 |
| Pre-treatment | THC (0, 5 mg/kg, i.p.) | N/A | THC (0-20 mg/kg, i.p.) | PG, CBD 400 mg/mL | Air, PG, THC 100 mg/mL | PG, CBD 400, THC 100 mg/mL | PG, CBD 400, THC 100 mg/mL | N/A | PG, CBD 100, CBD 400 mg/mL |

#### 2.4.3 CBD and THC Vapor Effects Under Progressive Ratio

Then rats were returned to the PR response contingency (0.15 mg/kg/infusion) for Sessions 65-69 and exposed to vehicle (PG) vapor, CBD vapor (400 mg/mL in the PG) or THC vapor (100 mg/mL in the PG) for 30 minutes immediately prior to IVSA sessions in a counter balanced order for Sessions 66-68. Thereafter the oxycodone dose was increased to 0.30 mg/kg/infusion for Sessions 70-74 and the PG, CBD and THC pre-exposure conditions repeated in a counterbalanced order in Sessions 71-73. For Sessions 75-77 the dose of oxycodone (0.06, 0.15, 0.30 mg/kg/infusion) was altered in a counterbalanced order, with no pre-treatment. One individual’s catheter exhibited resistance to daily flushing in this experiment and was non-patent by the final experiment and was therefore excluded from analysis. Rats completed PR sessions with no pre-treatment at the 0.03 mg/kg/inf dose for Sessions 78-83, and at the 0.06 mg/kg/infusion dose for Session 84.

#### 2.4.4 CBD and THC Vapor Effects Under Fixed-Ratio 1

Rats (N=9) were returned to FR1 in 3 h sessions with the oxycodone dose 0.06 mg/kg/infusion for Sessions 85-88. Rats were exposed to PG vapor, or CBD vapor (100, 400 mg/mL in the PG) for 30 minutes immediately prior to IVSA sessions in a counter balanced order for Sessions 86-88. One rat had a catheter that failed patency during this study and was therefore excluded.

### 2.5 Data Analysis

Infusions obtained, the percentage of responses directed at the drug-associated lever, total correct responses and breakpoints (for Progressive Ratio) were analyzed by ANOVA, or by mixed-effect models where there were missing values. Within-subjects factors of Dose or Pre-treatment were included where relevant. In our approach a single priming infusion is delivered if no responses have been made within 30 minutes of session initiation under FR (this is not included for PR). These are rare after the initial few sessions of acquisition and are thus reported qualitatively where present, but not formally analyzed. In all analyses, a criterion of P<0.05 was used to infer that a significant difference existed. Any significant main effects were followed with post-hoc analysis using Tukey (multi-level factors), Sidak (two-level factors) or Dunnett (to assess change relative to one of the Treatment Conditions.) correction. All analysis used Prism for Windows (v. 9.5.1; GraphPad Software, Inc, San Diego CA).

## 3 Results

### 3.1 THC Injection Increases Oxycodone Self-Administration in a PR Procedure

This experiment showed that THC (5 mg/kg, i.p.) increases breakpoints reached by female rats in the progressive ratio procedure (**Figure 1**), confirmed by a significant effect of pre-session Treatment Condition [F (2, 22) = 3.86; P<0.05]. One subject did not complete the 0.15 mg/kg with THC condition, thus the analysis was mixed effects.

**Figure 1:**
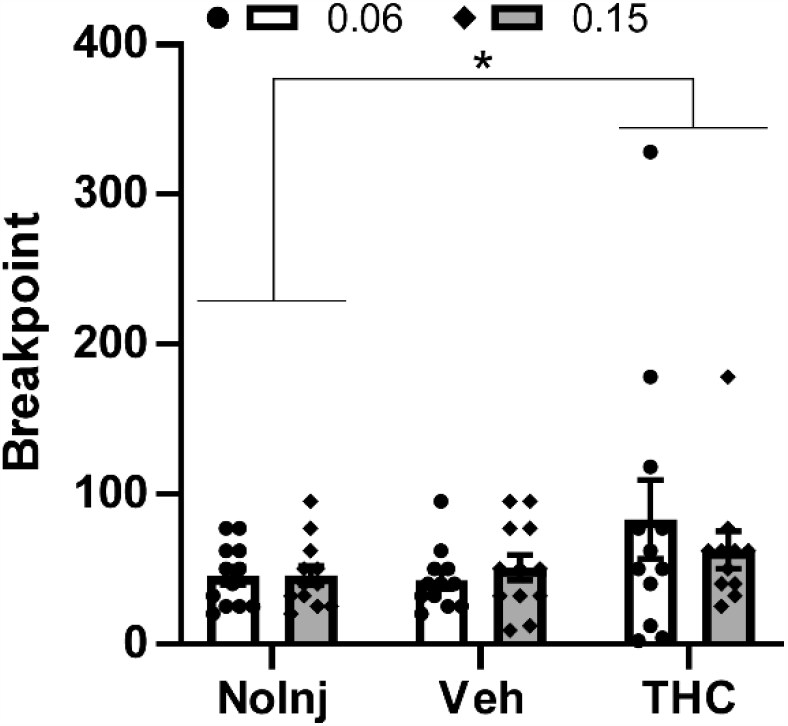
Mean (N=11-12 female rats; ±SEM) and individual breakpoints reached in the PR procedure following injection with Vehicle or THC (5 mg/kg, i.p.) in a counter-balanced order, followed by a no-injection day. The effect on self-administration of the oxycodone 0.15 mg/kg/infusion dose was evaluated first and the 0.06 mg/kg/infusion dose evaluated the following week. A significant difference between pre-treatment conditions, across oxycodone dose, is indicated with *.

The impact of THC was statistically indistinguishable across oxycodone unit doses (0.06, 0.15 mg/kg/infusion) as there was no significant effect of oxycodone Dose nor an interaction of Dose with Treatment Condition. The post-hoc Dunnett test of the marginal mean confirmed significantly lower breakpoints were reached in the No-Injection condition compared with the THC condition, across oxycodone Dose.

### 3.2 Oxycodone Self-Administration is Dose Dependent in a FR Procedure

The lack of difference between breakpoints generated under the different oxycodone unit doses prompted a comparison under FR1 conditions (3 h sessions to match the maximum duration of the PR sessions), to determine if these were functionally different doses in IVSA. This demonstrated that the 0.15 mg/kg/infusion dose resulted in fewer infusions obtained [Main Effect: F (2, 22) = 19.35; P<0.0001; **Figure 2A**], and generated fewer responses on the drug-associated manipulandum under the Time Out [Main Effect: F (2, 22) = 21.56; P<0.0001; **Figure 2C**], compared with the lower unit dose or vehicle. There was also a significant reduction in responses on the alternate manipulandum during the time-out [Main Effect: F (2, 22) = 3.92; P<0.05; **Figure 2D**] compared with the vehicle session.

**Figure 2:**
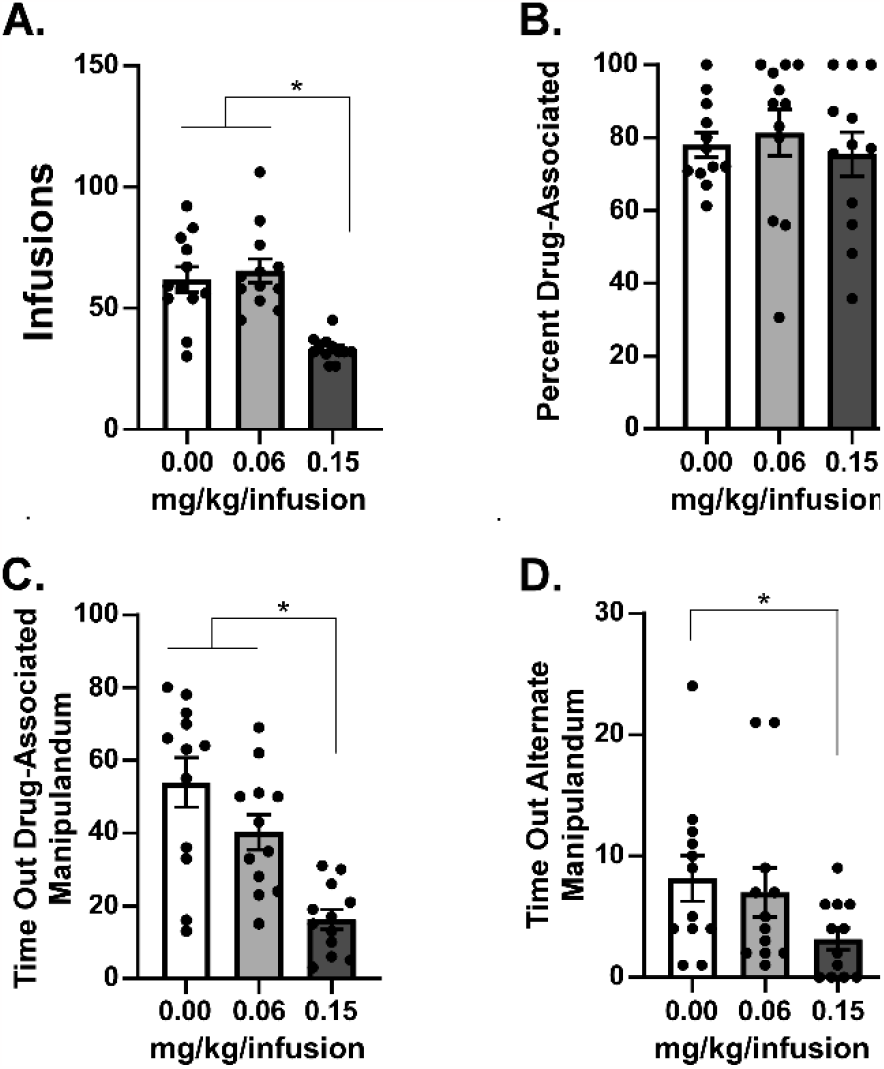
Mean (N=12; ±SEM) A) Infusions, B) Percent Drug-associated responses, C) responses on the drug-associated manipulandum during the time out and D) responses on the alternate manipulandum during the time out for self-administration of oxycodone in 3 h sessions under a FR1 response contingency. Tukey post-hoc test. A significant difference between pre-treatment conditions is indicated with *.

### 3.3 Cannabidiol and THC Decrease Oxycodone Self-Administration in a FR Procedure

CBD was delivered by vapor inhalation because in our hands it produces an involuntary responses (lowering of body temperature) in rats (Javadi-Paydar et al., 2019; Javadi-Paydar et al., 2018) whereas the injection of CBD at 20-60 mg/kg, i.p., has no effect (Taffe et al., 2015).

The impact of 30 min inhalation of vapor from CBD (400 mg/mL in the Propylene Glycol vehicle) prior to the IVSA session was then contrasted with the impact of vapor inhalation of THC (100 mg/mL in the PG). CBD inhalation significantly [F (2, 18) = 31.42; P<0.0001] reduced the number of infusions obtained in comparison with PG vapor inhalation or a session in which there was no exposure prior to the IVSA session (**Figure 3A**). Reponses on the drug-associated manipulandum were also significantly reduced compared with PG pre-treatment (F (2, 18) = 11.55; P<0.001; **Figure 3D**).

**Figure 3:**
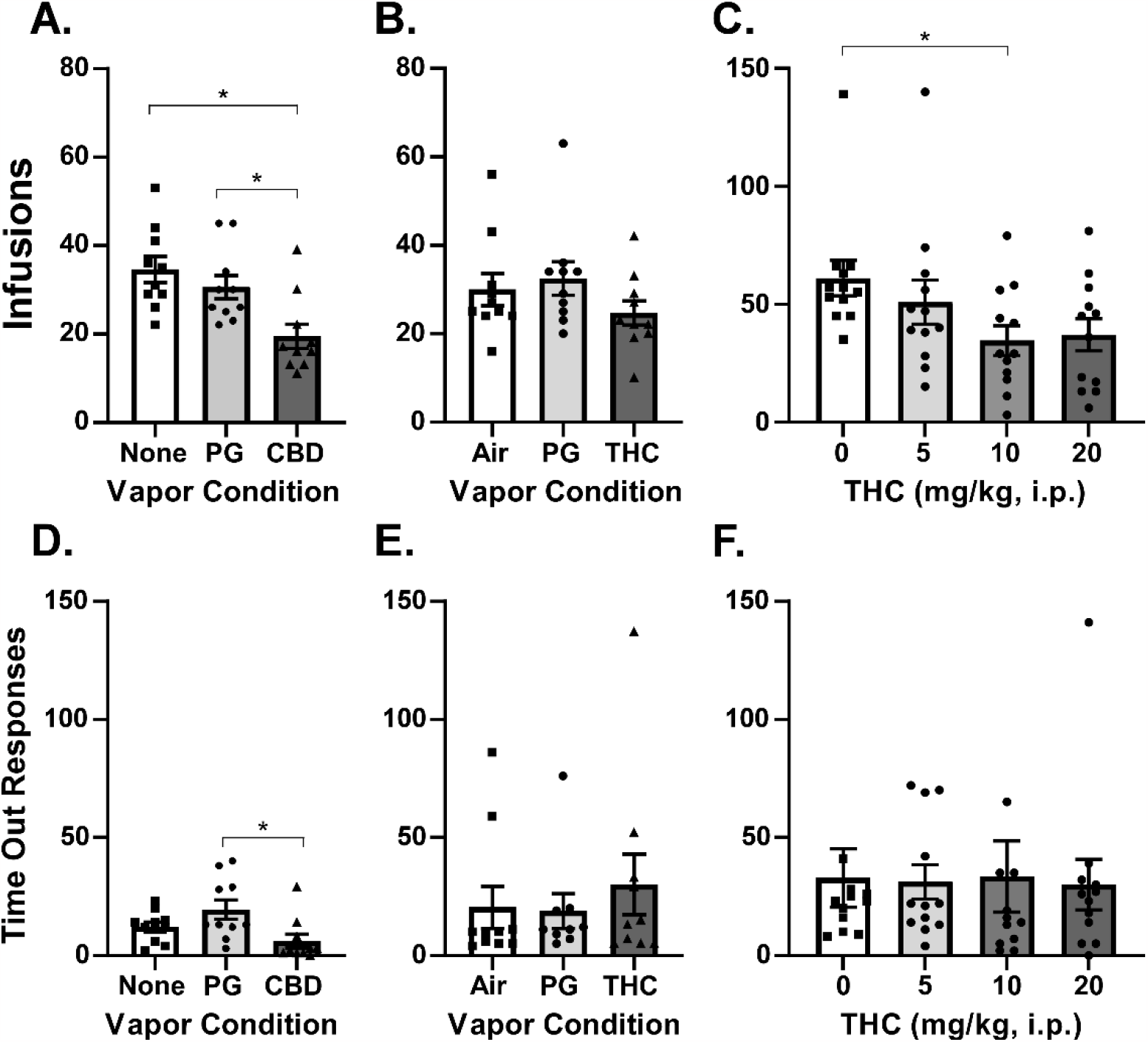
A) Mean (N=10; ±SEM) oxycodone (0.15 mg/kg) infusions obtained after exposure to vehicle (PG) or cannabidiol (CBD 400 mg/mL) vapor under a FR1 response contingency. B) Mean (N=10; ±SEM) oxycodone (0.15 mg/kg) infusions obtained after exposure to air, or PG or Δ^9^-tetrahydrocannabinol (THC; 100 mg/mL) vapor. C) Mean (N=12; ±SEM) oxycodone (0.06 mg/kg) infusions obtained after THC injection (0-20 mg/kg, i.p.) The corresponding responses on the drug-associated manipulandum is depicted in panels D, E and F. In panel F one individual responding 166 times in the vehicle condition and one individual responding 190 times after 10 mg/kg are omitted to facilitate scale comparison with the other panels. A significant difference between pre-treatment conditions is indicated with *.

THC vapor inhalation produced a small reduction in the infusions obtained (**Figure 3B**) which did not reach statistical significance (P=0.054). Considering that 5 mg/kg, i.p., may be a threshold dose (Nguyen et al., 2019), and perhaps subthreshold given prior repeated exposure of these rats to THC during acquisition or maintenance phases, we determined the effects of pretreatment with THC (5, 10, 20 mg/kg, i.p.) in a counter-balanced order. THC significantly [F (3, 33) = 2.93; P<0.05] reduced the infusions obtained and the Dunnett post-hoc test further confirmed that THC significantly reduced infusions obtained relative to vehicle injection at the 10 mg/kg, i.p., dose, but not at the 20 mg/kg, i.p. dose (P=0.065), (**Figure 3C**). There was a priming infusion delivered to one rat after 5 mg/kg and another rat after 10 mg/kg injection in this experiment.

### 3.4 THC, but not Cannabidiol, Inhalation Increases Oxycodone Self-Administration in a PR Procedure

It was a completely novel observation that CBD inhalation reduced oxycodone IVSA, so we then went on to further contrast the effects of inhaled PG, CBD and THC in a counter balanced order using a PR procedure. This was conducted first with the 0.15 mg/kg/infusion oxycodone dose and then with the 0.30 mg/kg dose (Wade et al., 2015), across sequential weeks.

THC inhalation increased drug seeking in the PR procedure (**Figure 4**). In contrast, CBD inhalation did not significantly alter oxycodone seeking relative to the PG inhalation condition. The analysis confirmed there were significant effects of vapor inhalation condition on Breakpoint [F (2, 32) = 5.36; P<0.01], Total Correct Responses [F (2, 32) = 5.59; P<0.01], Infusions [F (2, 32) = 4.47; P<0.05], and Time-Out Correct (responses on the drug-associated manipulandum) responses [F (2, 32) = 3.42; P<0.05]. The Tukey post-hoc tests of the marginal means confirmed that, across oxycodone dose, more infusions were obtained (**Figure 4C**) and more Time-Out correct responses were made (**Figure 4D**) after THC inhalation compared with CBD inhalation.

**Figure 4:**
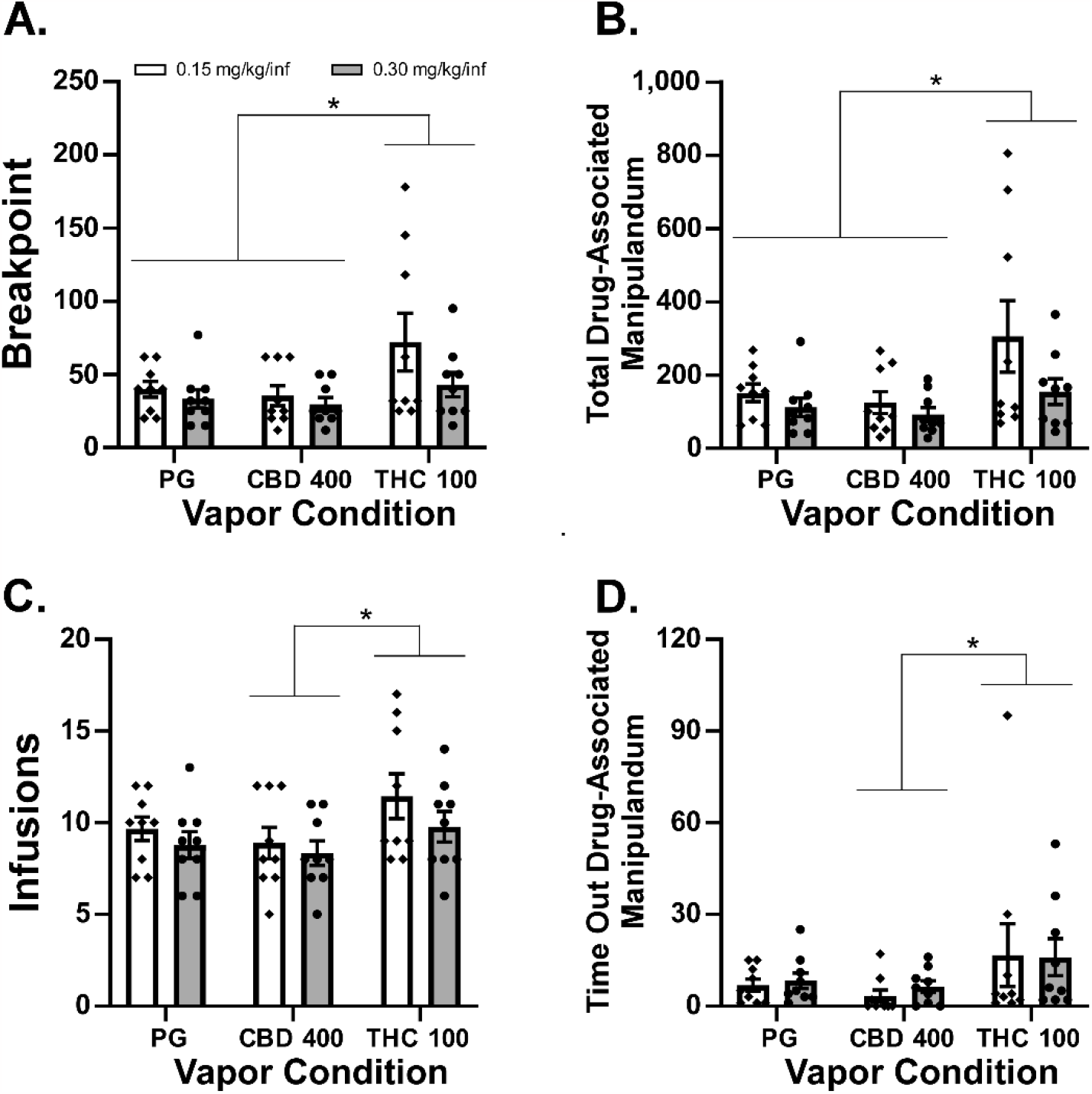
Mean A) Breakpoint, B) Total responses on the drug-associated manipulandum, C) Infusions of oxycodone (0.15, 0.30 mg/kg) obtained and D) Responses on the drug-associated manipulandum during the time-out interval during self-administration under a Progressive-Ratio response contingency. Significant differences associated with pre-treatment condition, across oxycodone dose, are indicated with *.

In addition, rats reached higher breakpoints (**Figure 4A**) and emitted more total correct responses (**Figure 4B**) after THC vapor inhalation compared with either PG or CBD vapor inhalation.

### 3.5 Cannabidiol Inhalation Decreases Oxycodone Self-Administration in a FR Procedure

#### FR1 oxycodone dose-response

We re-determined the impact of a range of oxycodone doses (0.06-0.3 mg/kg/infusion) under FR1 because the oxycodone dose did not alter self-administration of oxycodone under a PR procedure. The one-way ANOVA confirmed a significant impact of oxycodone Dose [F (2, 14) = 60.19; P<0.0001] on infusions obtained and the Tukey post hoc further confirmed significant differences between all doses (**Figure 5A**).

**Figure 5:**
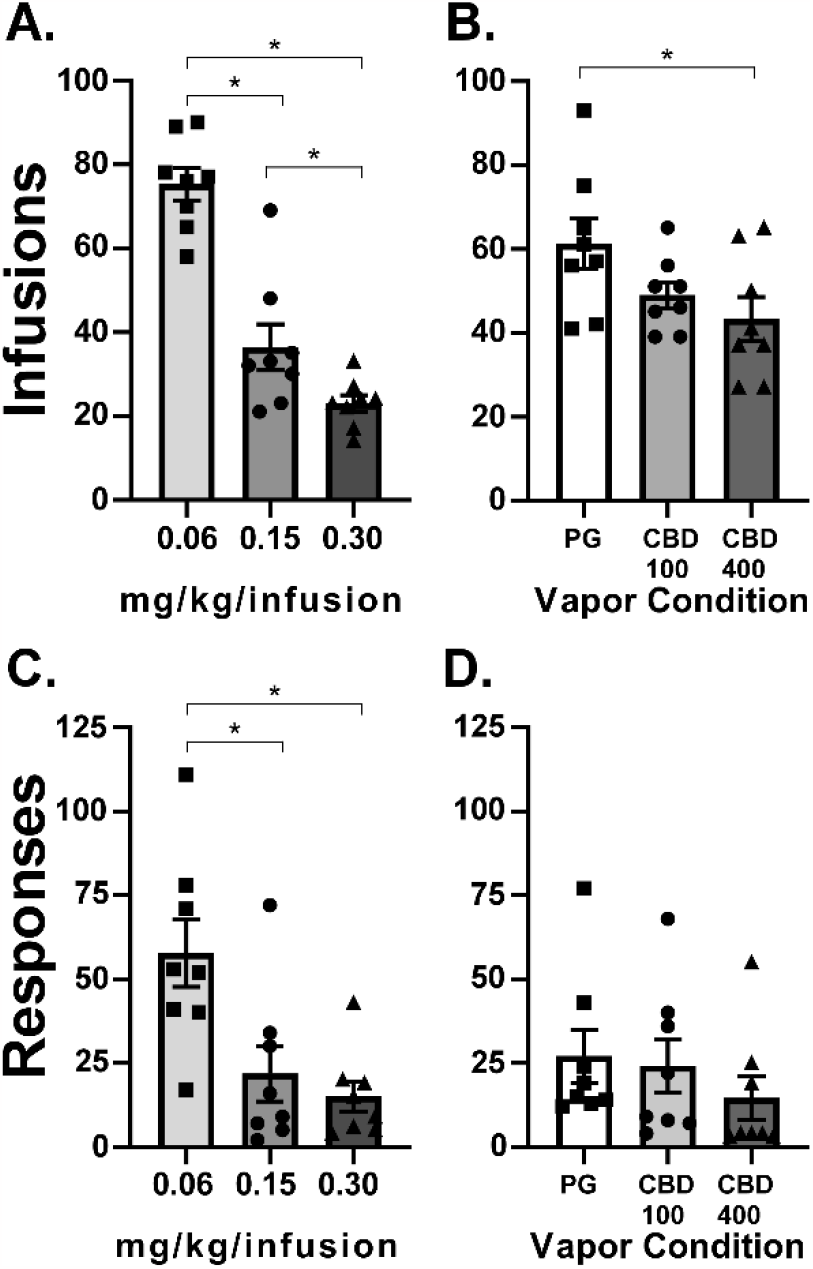
A) Mean (N=8; ±SEM) oxycodone infusions (0.06-0.30 mg/kg) obtained in 3 h sessions under an FR1 response contingency. B) Mean (N=8; ±SEM) oxycodone (0.06 mg/kg) infusions obtained in 3 h sessions under a FR1 response contingency after exposure to vehicle (PG) or cannabidiol (CBD 100 and 400 mg/mL) vapor.

#### FR1 CBD dose-response

Because CBD did not alter oxycodone IVSA under a PR procedure, we re-determined the impact of CBD vapor under FR1 to rule out a potential development of tolerance to the CBD. For this we tested two CBD vapor doses (100 and 400 mg/mL) and the 0.06 mg/kg/infusion dose of oxycodone. The one-way ANOVA confirmed a significant impact of Pre-Treatment condition [F (2, 14) = 5.79; P<0.05] on infusions obtained and the Tukey post-hoc further confirmed significantly fewer infusions were obtained after CBD 400 exposure compared to PG exposure (**Figure 5B**).

## 4 Discussion

This study replicates and generalizes our prior work by showing that Δ^9^-tetrahydrocannabinol (THC) *increases* female rats’ oxycodone seeking under a Progressive Ratio schedule, while it *reduces* oxycodone self-administration under a Fixed Ratio 1 (FR1) contingency. Our previous study was conducted in male rats and while several conditions of oxycodone IVSA were evaluated, it was all conducted with a FR1 schedule of reinforcement (Nguyen et al., 2019). In that prior study, however, THC did increase breakpoints in a PR procedure when the animals were switched to the IVSA of *heroin* (0.006, 0.06 mg/kg/infusion doses) and the present study found that the effect of FR vs PR reinforcement schedule on the impact of THC extended to *oxycodone* IVSA in female rats. The increased behavioral output (e.g., total drug-associated responses) in the PR procedure associated with THC (**Figure 1, 4**) is also inconsistent with an inference that reductions in drug seeking in the FR1 procedure are due to general sedating properties of the THC.

Although inhaled THC did not significantly reduce oxycodone IVSA (**Figure 3B**) as it did in our prior study, this is almost certainly a matter of THC dose, possibly related to tolerance created by the repeated THC pre-treatments during the prior study in these rats. This interpretation is supported by the injection experiments (**Figure 3C**), which established that THC delivered intraperitoneally reduces oxycodone IVSA under a FR1 contingency in the female rats, just as it did with males in our prior study (Nguyen et al., 2019). Thus, the lack of effect of the inhaled THC was likely due to dose, since the THC injection study showed that 10 mg/kg, but not 5 mg/kg, was able to reduce oxycodone IVSA.

It was somewhat unexpected that rats reached similar breakpoints for 0.06 and 0.15 mg/kg/infusion in the PR under control conditions (PG here, also true in the no-treatment acclimation sessions) given an apparent difference across this dose range that was observed in prior work with male rats trained in sessions of 1 h or 12 h duration (Nguyen et al., 2021; Nguyen et al., 2018; Wade et al., 2015). In particular, Wade and colleagues showed that the largest difference was between the 0.06 and 0.15 mg/kg/infusion doses in the animals trained in 12 h sessions and between the 0.15 and 0.30 mg/kg/infusion doses in the animals trained in 1 h sessions. In this study rats did, however, exhibit differential intake for 0.06, 0.15 and 0.3 mg/kg/infusions under the FR1 response contingency (**Figure 5A**), demonstrating the expected behavioral compensation. In our prior study, male rats self-administered more infusions at the 0.06 mg/kg dose compared with the 0.15 mg/kg dose under an FR1 response contingency, and this did not interact with the effects of THC (Nguyen et al., 2019). However, there was no effect of *heroin* unit dose (0.006 vs 0.06 mg/kg/infusion) in a PR procedure, nor any interaction of the effect of THC with heroin unit dose, in that study. Therefore, the prior results were quite similar to the impact of unit dose of oxycodone across FR1 and PR procedures in the current study.

One of the strengths of this study was that animals were originally trained in extended access (6 h) sessions, examined for this study in 3 h sessions under both FR1 and PR response contingencies and exhibited a high degree of behavioral stability across an extended number of sessions. This enhances comparison of findings across the experiments. In general, this confirms ongoing work from multiple laboratories showing that oxycodone is a typical opioid reinforcer in rat intravenous self-administration, and that stable self-administration patterns are produced in short-access IVSA in male and female rats (Mavrikaki et al., 2017; Pravetoni et al., 2014). Similarly, behavioral escalation, and disruptions of brain reward systems have been observed with extended access IVSA (Blackwood et al., 2019a; Blackwood et al., 2019b; Matzeu and Martin-Fardon, 2020; Nguyen et al., 2021; Nguyen et al., 2017; Wade et al., 2015).

In contrast with THC, the impact of CBD was to reduce oxycodone seeking under FR1 and to leave it unaffected under the PR1 contingency. CBD has been previously shown to have anti-drug-seeking effects in rats (Gonzalez-Cuevas et al., 2018) when delivered by the transdermal route and to diminish withdrawal associated with chronic nicotine exposure (Smith et al., 2021) when injected repeatedly with the nicotine. CBD i.p. also reduced cocaine IVSA in rats under FR1 and PR schedules of reinforcement with the effects significant in lower unit doses of cocaine but not with higher unit doses of cocaine (Galaj et al., 2020). These effects were mediated by CB2, TRPV1 and 5HT1A receptors but not by CB1, GPR55 or MOR receptors. Effects of inhaled CBD in the present study were therefore consistent with results of Gonzalez-Cuevas and colleagues (2018), since oxycodone infusions were reduced in the FR1 study. CBD also failed to affect cocaine IVSA under a PR schedule in rats in prior study (Mahmud et al., 2017). This latter finding in the present study functions as a negative control to further emphasize the impact of THC is to increase, not decrease, the rewarding value of a unit dose of oxycodone.

Interestingly, the inhalation of vapor from cannabis extracts with high CBD content (64.2% CBD and 7.1% THC) decreased fentanyl IVSA under a FR, but not a PR, schedule in rats with and without induced neuropathic pain (Rivera-Garcia et al., 2023). Interactive effects of CBD administered with THC remain unclear in rodent models (Boggs et al., 2018), however this result implies that the impact of the CBD overcame the impact of the THC in that study, rather than enhancing the impact of THC, e.g., via metabolic inhibition (Varvel et al., 2006).

In conclusion these studies provided additional evidence that THC reduces oxycodone seeking under easy-access conditions because it enhances the rewarding value of a unit dose of oxycodone. When access requires more work in a PR procedure, THC increases oxycodone seeking responding. In contrast, the effect of CBD was to decrease oxycodone self-administration in easy access conditions but to have no effect in harder access conditions. This cautions that cannabis is not necessarily reducing addiction

## Acknowledgements

These studies were supported by the UCSD Center for Medicinal Cannabis Research (P64-04-002 originally awarded to JDN), UCSD Medical Scientist Training Program Summer Undergraduate Research Fellowship (CY), the UCSD PATHS Scholar Program (CY), UCSD Chancellor’s Post-doctoral Fellowship (AG), UCSD IRACDA (K12 GM068524; AG), the Tobacco Related Disease Research Program (T31IP1832; MAT) and the NIH (K99 DA047413; JDN).

